# Feature extraction through Wavelet decomposition for automatic detection of landmarks on Cephalogram Images

**DOI:** 10.1101/125286

**Authors:** Srikanth Vasamsetti, Viren Sardana, Prasoon Kumar, H.K Sardana

## Abstract

Manual cephalogram marking has long way from marking on tracing sheet to availability of commercial software's for cephalometric analysis. With the effort involved in manual marking and time consumption, it becomes imperative for modern science to envisage algorithms which could automatically locate landmarks on the cephalogram images and perform various analysis. In this work, we herby propose a wavelet transform based feature extraction algorithm for detection of landmark on cephalogram images. 15 landmarks were detected on the images using wavelet transform and all landmarks were detected within the acceptable accuracy limits. This algorithm may have a promising approach in detection of further anatomical landmarks automatically and analysis and thus may help orthodontic practioners in better and faster treatment planning.

## I. INTRODUCTION

Cephalometrics is a technique which assists orthodontic surgeons to evaluate head measurements through oriented radiographs. It generates qualitative description of the craniofacial deformities and presents the complexities of human head into simple geometric scheme comprising of distances, ratios and proportions, angles for proper assessment and comparison with normal human cranium [1]. Orthodontists use cephalometric analysis of lateral and posterior skull X-Rays for craniofacial growth assessment, diagnosis of facial deformities, treatment planning and craniofacial surgery in malocclusions. The concept of cephalometry was introduced for the first time in year 1931 by Broadbent [2]. The current practice in the field of orthodontics involves the use of 2-D x-ray radiograph images. They overlay a tracing sheet on cephalogram film for locating the anatomical landmarks, drawing lines and angle with scale and protractor respectively through their own experience and expertise. As the aforementioned method is needed to be carried out for every patient, it is subjected to human error and is tedious, time consuming and subjective in nature, eventually affecting proper treatment planning. Thus automating landmark detection with analysis through image processing techniques in engineering can provide assistance to orthodontic surgeons with rapid and increased fidelity in treatment planning. The automated cephalometry will eliminate inter and intra subject variability in addition to improved efficiency and accuracy.

Research in automatic landmark detection through image processing can be classified in four stages of development. Landmark detection on cephalogram was automated and was primarily dependent on edge detection techniques as suggested by Levy-Mandel et al. [3], Jackson et al. [4] and Cohen et al. [5]. The major limitations associated with aforementioned technique are the practical constraints in obtaining high quality cephalograms and inability to track the landmarks away from edges as detection is dependent on the quality of the edges in the image and geometric definition of a particular landmark. The second generation researchers and scientists used mathematical and statistical models to optimize the search area over the cephalogram and thereafter implemented the shape matching techniques for accurate positioning of landmarks. Cardilo and Sid-Ahmed [6], Grau et al [7] demonstrated the application of above method on detection of landmarks within an acceptable error limit. The errors in detection are major limitation. Aforementioned limitations led to the deployment of neural networks and fuzzy inference system for detection of landmarks by the third generation researchers. Chen et al [8], Chakrabarty et al. [9], Innes et al. [10] used the tools of soft computing for automating the landmark detection on cephalogram. In spite of them being successful in designing methods for automatic landmark detection, but due to greater human intervention in parameterization, the very idea was abandoned. The debacle of the aforementioned method caused the inception of fourth generation researchers who utilised the combination of above methods. Yue et al. [11], Kafieh et al. [12], HadisMohseni and Shohrehkasaei[13], I.EI-Fegh et al [14] have demonstrated the potential of combination of various above methods for effective localization of landmarks automatically. Earlier work carried in our lab by Jain et al. [15] suggested the promising nature of the template matching towards automatic cephalometric analysis.

Although numerous methods for cephalometric automation have been enumerated but still noise, rough texture, occlusion, illumination changes with weak edges has been stumbling block in developing a generalized method for automatic landmark detection. As the cephalograms have definitive features, so feature extraction technique which overcomes above limitation could be a promising method for automating landmark detection. The feature extraction can be carried out through multilevel resolution of images into spatial and frequency domain through various transforms. The literature suggests that the application of wavelet based method for feature extraction in a number of applications [15,16]. The wavelet transform has been one of the premier methods which can lead to multiresolution decomposition to capture the features of an image at all scales. This generates the information at various scales which can be manipulated to extract information regarding the location of various features on the image. In this paper, we have demonstrated the use of one of the variant of discrete wavelet transform for anatomical feature extraction on the cephalogram and thereby detection of landmarks.

## II. MATERIALS AND METHODS

### A. Materials

The proposed algorithm was developed using MATLAB R2011a and was tested on a set of 20 lateral cephalogram images provide by All India Institute of Medical Sciences, New Delhi. Result verification was performed using data obtained by marking of the landmarks on all the images by a practicing orthodontic specialist.

### B. Methods

In the present method, we have developed a wavelet decomposition coupled optimized template matching algorithm for localization of anatomical landmarks on the cephalograms. The algorithm comprises of three major steps. First step involves the search of the cephalostat region on cephalogram to mark it as a reference point for the search of other regions. The cephalostat region of cephalogram is chosen in liu of its distinct foreground and background, thereby enabling its quick and accurate detection. The searching of region of interest is carried out through feature extraction method. The two tier feature identification method is employed, one for the coarse features and other for the fine features. The fine search is carried out in the region delimited by course feature region. The step size during coarse search is greater than fine search aiding in quick search without compromise on the accuracy of the results. The number of template used to locate a land mark region precisely is dependent on the landmark position. For instance, as due to huge variability in the incisor region, the number of template used in its detection is greater than the template used for detection of tip of nose.

A cephalogram on which landmarks are to be located is taken and divided into four quadrants followed by the step of template image of the cephalostat as mentioned before is used and searched in the upper right quadrant of the image. This is followed by the template image and the region of search in the reference image being subjected to first level decomposition by haar wavelet. Thereafter the first level decomposed image is taken and its approximate coefficients are further subjected to second level decomposition to generate the horizontal, vertical, diagonal and approximate coefficients of second level decomposed image.

**Fig 1.**
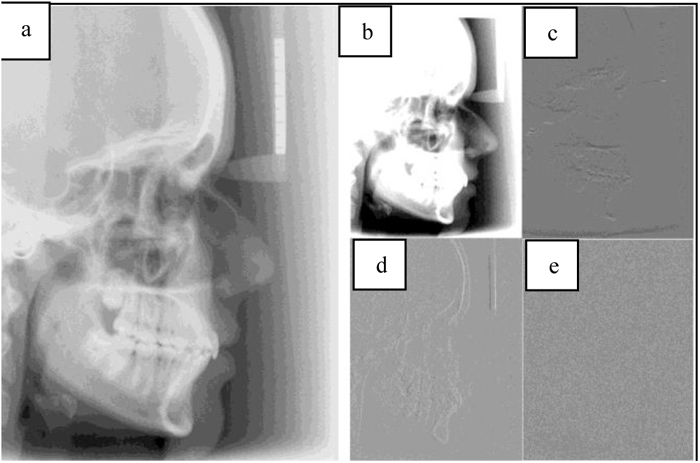
One level decomposition of cephalometric image by using Haar wavelet a) Test image, b) Low frequency sub band image, c) Horizontal high frequency sub band image, d) Vertical high frequency sub band image, e) Diagonal high frequency sub band image

Then a virtual image is generated by averaging the aforementioned second level horizontal, vertical and diagonal coefficients for the template as well as the searched region of the reference image. The virtual image generated from the test image and reference image are taken as input images for determination of correlation coefficients between the two images. Correlation coefficient is calculated using equation 1.

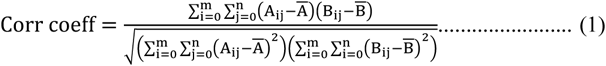
where, A, B are input images, 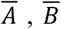 represents the average values of A and B and m, n are the rows and columns of the input image.

During the search process, the region which shows the maximum correlation coefficient value is selected for the further processing to locate the landmarks. This method is employed during every searching step, irrespective of the nature of search: course or fine for locating the precise region of landmarks. Schematic bock diagram of the approach used is depicted in Figure 2. Figure 1 depicts that sample has been decomposed into four sub images using haar wavelet transform. The high frequency information in the horizontal, vertical and diagonal directions and low frequency information have been extracted and stored in each sub image. Figure 3 depicts low frequency sub band image and depicts combination of horizontal, vertical and diagonal sub band images.

**Fig 2.**
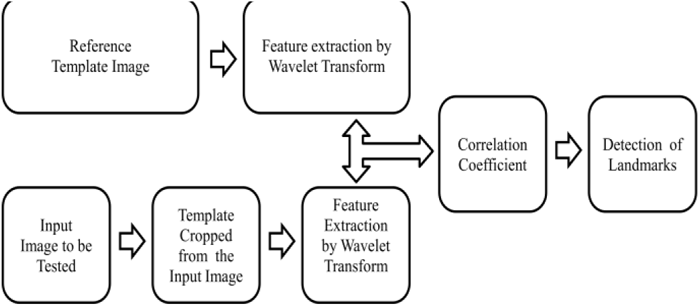
Schematic for the proposed Algorithm

**Fig 3.**
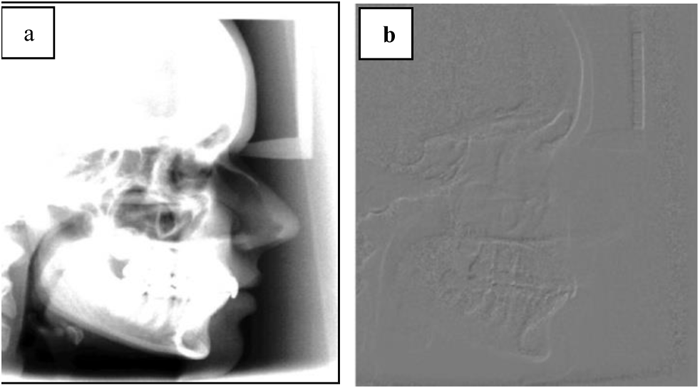
a) low frequency sub image b) Recombined high frequency sub image

The region obtained through aforementioned methodology is used as the region for searching the landmarks required for cephalometric analysis. Depending upon the location of the landmarks, two approaches have been implemented to determine the exact landmark position on the cephalogram. The landmark localization carried out on the basis geometric definition or on prior knowledge of its whereabouts on the edges of regions in cephalograms. Once the smallest region with particular landmarks is determined, and then according to the landmark, one of the aforementioned approaches is adopted. The former approach of geometric definition has been used as measure to locate landmarks like Sella, Orbitale. For example, the Sella is located on cephalogram through its geometric definition which states that Sella is the point which is the mid-point of the hypophyseal fossa. Once the tertiary or secondary region within which landmark localized has been finalised, then geometric feature and its relationship with the given landmark is being used as measure to determine landmarks on the cephalogram. The latter approach of determination of landmarks based on prior knowledge is used to determine landmarks like B-point, Nasion, Soft Nasion, Pronasale, Subnasale, Labrale inferius, Labrale superius, Soft B-point, Pogonion, Menton, Gnathion and Soft Pogonion. Once the tertiary or secondary region within which landmark localized has been finalized, the edge detection is performed to obtain the binary image. For instance, the binary image after edge detection in cephalogram indicates an arc in B-point region. The determination of landmarks on the cephalogram is carried out based on its location on the edge after binarization. Considering arc to be a part of a complete circle, the deepest point from the left of the image is taken as B-point. Similar approach has been used to determine the other landmark is this category. As the images suffer from the variation in luminosity, the image quality affects the edge detection methodology. So, the image enhancement technique like histogram equalization was carried out before edge detection. Thereafter canny edge detection was employed with optimized parameters. Eventually position of the landmark, depending upon their specific location on the edges with respect to boundaries of images, was located on the cephalogram. For example, Nasion is the most anterior point of the nasofrontal in the median plane and is present in a region of the image where the machine tripod touches the nasofrontal curve. As there is existence of soft and hard nasion in aforementioned region, so care must be taken while locating a particular land marks.

The quality of the cephalogram is affected due to variations in brightness and contrast between inter-images and intra-images. Due to region wise variations in the images intensity, block processing of image were carried out before localization of landmarks. The different regions of the block were enhanced by adaptive histogram equalization, thereby minimization of error during detection due to illumination difference. Thereafter canny edge detection was carried out on these enhanced images with optimized parameters. The automatic thresh holding was done by Otsu’s Method [16] for binarization and edge detection. Then images were subjected to morphological operations like repeated dilation and erosion for improvement on the quality of the edges detected on the cephalogram. The prior knowledge of landmarks on the edges so detected has been used as measure to locate the land marks.

## III. RESULTS AND DISCUSSION

The data obtained from 20 images marked by an orthodontic specialist for localization of 15 anatomical landmarks of which 7 were bony landmarks and 8 were soft tissue were used for verification of the proposed algorithm's accuracy. From the previous literature, inter subject variability was such that landmarks detected within 2mm were termed accurate and acceptable within 5mm [11, 17, 18]. Considering the landmark point marked by orthodontic specialist on cephalogram as reference, we have calculated the mean error for each landmark in x and y directions and mean error calculated using equation 2.

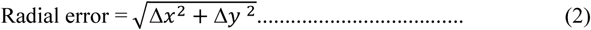
When we compared the results of landmarks marked by orthodontic specialist with our algorithm detected landmarks, it was observed that 5 landmarks out of 15 (3 bony points and 2 soft tissue points) were on the limit of the accurate limitation value of 2 mm while the 10 points were under 2 mm for mean error values, while for the individual average *x* and *y* differences, all values were under 2 mm, hence proving the accuracy in automatic localization of landmarks on cephalogram images. Figure 4 demonstrates the landmarks detected using the proposed wavelet transform algorithm.

**TABLE 1.** Localisation errors of landmarks

| S.No | Anatomical Landmarks | Avg. $\Delta x$ (mm) | Avg. $\Delta y$ (mm) | Radial error (mm) |
| --- | --- | --- | --- | --- |
| 1 | Sella | 0.63 | 0.35 | 0.72 |
| 2 | Nasion | 1.22 | 1.61 | 2.02 |
| 3 | Orbitale | 1.32 | 1.70 | 2.16 |
| 4 | B-point | 0.84 | 1.84 | 2.03 |
| 5 | Pogonion | 0.31 | 0.96 | 1.00 |
| 6 | Menton | 0.69 | 0.19 | 0.72 |
| 7 | Gnathion | 0.79 | 0.59 | 0.99 |
| 8 | Pronasale | 0.51 | 1.35 | 1.44 |
| 9 | Subnasale | 1.16 | 1.87 | 2.20 |
| 10 | Soft A-point | 0.45 | 1.66 | 1.72 |
| 11 | Labrale superius | 0.96 | 1.13 | 1.48 |
| 12 | Labrale inferius | 0.67 | 1.76 | 1.88 |
| 13 | Soft B-point | 0.37 | 1.63 | 1.67 |
| 14 | Soft Pogonion | 0.94 | 1.93 | 2.15 |
| 15 | Soft Tissue Nasion | 0.79 | 1.25 | 1.48 |

During wavelet decomposition, the signals corresponding to edges are extracted in terms of horizontal, vertical and diagonal coefficients, thereby enabling the easy detection of landmarks on edges. The landmarks not lying on edges are present in approximate coefficient details which represent a weak signal. The extraction of features in weaker signals is difficult and need to filter for getting meaningful signal. This suggests that the quality of image affects the landmark accuracy. For the detection of landmarks in the region with feeble signal, it is needed to be development of filters to extract features corresponding to weak signals. The values obtained for individual x an y differences and mean error have been depicted in table 1 for all the landmarks.

## IV. CONCLUSION

The automated cephalometric analysis has been one of the desired tools aspired in the field of dentistry. This would not only reduce inter subject variability during landmarks detection but also reduce time and resources without affecting the treatment planning. In this paper, we have developed a feature extraction algorithm through wavelet which is capable of landmark detection on digital Cephalogram. We applied our method to 20 cephalogram images for automatic detection of anatomical landmarks and successful detection was achieved for 7 bony and 8 soft tissue landmarks within the acceptable accuracy range of 2 mm. Our method is robust to anatomical variations, intensity modulation but affected by edge features due to implementation of wavelet decomposition followed template matching method. In future, we need to develop filters to extract features represented by weaker signals. Further promises in the algorithm lie in its extrapolation to detect more landmarks on higher number of cephalograms which will be helpful in generating complete understanding of the cephalometric analysis.

**Fig 4.**
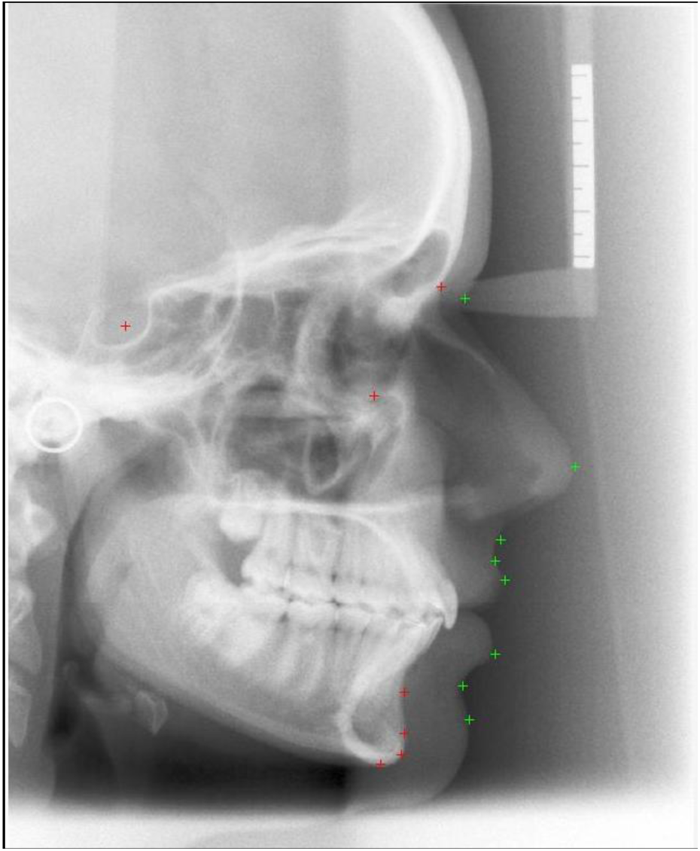
Landmarks detected on a cephalogram using wavelet transform algorithm

## ACKNOWLEDGEMENT

Authors would like to thanks All India Institute of Medical Sciences, New Delhi for cephalogram data and Dr Poonam for marking of Image data, Mamta Sharma for her assistant to orthodontists for locating landmarks on cephalometric images, Aparna Akula and Satish Kumar for their valuable suggestions in research work.

